# Migration and interaction in a contact zone: mtDNA variation among Bantu-speakers in southern Africa

**DOI:** 10.1101/002808

**Authors:** Chiara Barbieri, Mário Vicente, Sandra Oliveira, Koen Bostoen, Jorge Rocha, Mark Stoneking, Brigitte Pakendorf

**Affiliations:** Department of Evolutionary Genetics, MPI for Evolutionary Anthropology, Leipzig, Germany, 04103; Department of Biological, Geological and Environmental Sciences, Laboratory of Molecular Anthropology, University of Bologna, Bologna, Italy, 40126; CIBIO, Centro de Investigação em Biodiversidade e Recursos Genéticos da Universidade do Porto, Vairão, 4485-661, Portugal; STAB VIDA, Investigação e Serviços em Ciências Biológicas, Lda, Oeiras 2780-182, Portugal; Division of Biological Anthropology, University of Cambridge, Cambridge CB2 3QG, UK (current affiliation); Departamento de Biologia, Faculdade de Ciências da Universidade do Porto, Porto, 4169-007, Portugal; Department of African Languages and Cultures, Ghent University, KongoKing Research Group, B-9000 Ghent, Belgium; Université libre de Bruxelles, Faculté de Philosophie et Lettres, B-1050 Brussels, Belgium; Laboratoire Dynamique du Langage, UMR5596, CNRS and Université Lyon Lumière 2, Lyon, France, 69007

## Abstract

Bantu speech communities expanded over large parts of sub-Saharan Africa within the last 4000-5000 years, reaching different parts of southern Africa 1200-2000 years ago. The Bantu languages subdivide in several major branches, with languages belonging to the Eastern and Western Bantu branches spreading over large parts of Central, Eastern, and Southern Africa. There is still debate whether this linguistic divide is correlated with a genetic distinction between Eastern and Western Bantu speakers. During their expansion, Bantu speakers would have come into contact with diverse local populations, such as the Khoisan hunter-gatherers and pastoralists of southern Africa, with whom they may have intermarried. In this study, we analyze complete mtDNA genome sequences from over 900 Bantu-speaking individuals from Angola, Zambia, Namibia, and Botswana to investigate the demographic processes at play during the last stages of the Bantu expansion. Our results show that most of these Bantu-speaking populations are genetically very homogenous, with no genetic division between speakers of Eastern and Western Bantu languages. Most of the mtDNA diversity in our dataset is due to different degrees of admixture with autochthonous populations. Only the pastoralist Himba and Herero stand out due to high frequencies of particular L3f and L3d lineages; the latter are also found in the neighboring Damara, who speak a Khoisan language and were foragers and small-stock herders. In contrast, the close cultural and linguistic relatives of the Herero and Himba, the Kuvale, are genetically similar to other Bantu-speakers. Nevertheless, as demonstrated by resampling tests, the genetic divergence of Herero, Himba, and Kuvale is compatible with a common shared ancestry with high levels of drift and differential female admixture with local pre-Bantu populations.

## INTRODUCTION

Bantu languages started to diffuse from their homeland in the Grassfields of Cameroon around 4,000-5,000 years ago, reaching the southernmost areas of the continent in only a few thousand years [1–5]. This spread, strongly associated in its later phases with the diffusion of technological advances related to metallurgy and an agricultural lifestyle, was probably the result of a long-distance migration of people who partially replaced the local forager and pastoralist populations, or intermixed with them [2,6,7]. From a linguistic perspective, the genealogical unity of the Bantu family is certain, even though its boundary with other branches of the Niger-Congo phylum is not clear-cut and the internal classification and distinction between languages and dialects is highly debated [4,8]. The region close to the putative homeland represents the highest linguistic diversity. The first Bantu branches to split off, such as Mbam-Bubi and North-West Bantu, are confined to Cameroon and immediately neighboring regions [9]. The remainder of the Bantu languages predominantly belongs to two major branches, namely Eastern Bantu and Western Bantu, which are further divided in several subgroups. Although a recent investigation finds a distinct trace of the eastern route of the Bantu migration in Y-chromosomal variation [10], other molecular anthropological studies fail to find evidence for a genetic differentiation of the populations speaking Western and Eastern Bantu languages [11,12].

Southern Africa represents the last phase of the Bantu expansion. Archaeological data reveal traces of an agricultural way of subsistence in Namibia, Zambia and Botswana around 2000-1200 years ago [1,13,14], which was preceded by a few centuries by an immigration of pastoralist cultures [15,16]. Thus, in these areas, the presumably Bantu-speaking agriculturalist immigrants would have met both populations of hunter-gatherers as well as pastoralists, whose descendants comprise the linguistically, culturally, and genetically diverse “Khoisan” populations [17,18].

The Bantu-speaking populations nowadays inhabiting southern Africa are quite diverse linguistically and culturally, comprising pastoralists, agro-pastoralists, and agriculturalists who speak languages belonging to several different subgroups of both Eastern and Western Bantu; these populations share the same territory and are often involved in trade. From a genetic perspective, these populations appear to be relatively homogenous, with little differences even among linguistically distinct populations [6,11,12,19]. The main genetic signal characterizing the people at the southernmost edges of the Bantu expansion is the degree of admixture with the autochthonous populations; this can be explicitly measured by the presence of the characteristic mtDNA haplogroups L0d and L0k and Y-chromosomal haplogroups A-M51, A-M23, and B-M112 [20–22]. Admixture with autochthonous peoples in Bantu-speaking populations is detectable predominantly in the maternal line, in accordance with sex-biased gene flow [20,23]. The level of admixture differs considerably among populations; in particular, substantial proportions of mtDNA haplogroups L0d and/or L0k are observed in the pastoralist Kuvale from southwestern Angola [21], in the Fwe of southwestern Zambia [20], and in the Zulu and Xhosa from South Africa [24]. In contrast, in populations from eastern Zambia, Zimbabwe, and Mozambique these characteristic autochthonous haplogroups are found at a frequency of at most 3% [19,25,26].

Among the culturally distinct populations in southern Africa are the Herero, Himba, and Kuvale from northern Namibia and southern Angola, who speak dialects of the same Bantu language and practice intensive semi-nomadic cattle pastoralism. The Herero and Himba appear genetically distinct from other Bantu-speaking populations of the area, including the culturally similar Kuvale [17,21,27,28]. Genetically, the closest relatives of the Herero and Himba are the Damara [17,18], hunter-gatherers and small stock herders who speak a Khoisan language of the Khoe-Kwadi family [29]. Intriguingly, the Herero were known as “Cattle Damara” and the Damara were referred to as “Berg Damara” in previous literature [29].

In this study we analyzed complete mtDNA genome sequences of 944 Bantu-speaking individuals from Angola, Zambia, Namibia, and Botswana to investigate the maternal genetic history of Bantu speakers of southern Africa. We also include 38 Damara mtDNA genome sequences to further investigate the close genetic relationship between the Herero, Himba, and Damara that emerged in previous research [18]. We focus on the following research questions: 1) does the linguistic division into Western and Eastern Bantu correlate with genetic divergence? 2) To what extent did the immigrating Bantu-speaking agriculturalists intermarry with autochthonous populations? 3) What factors can explain the genetic divergence between the culturally and linguistically closely related Himba, Herero, and Kuvale on the one hand, and the genetic proximity of the Himba and Herero to the culturally and linguistically very distinct Damara? Our results reveal a general homogeneity of the maternal lineages of Bantu speakers of Angola and Zambia and suggest different demographic histories for the Herero, Himba and Damara from Namibia as well as for Bantu-speaking populations of southern Botswana.

## MATERIALS AND METHODS

### Ethics statement

The collection of the samples was approved by the Ethics Committee of the University of Leipzig and the Research Ethics Committee of the University of Zambia. Published samples from Botswana and Namibia come from a sample collection authorized by the governments of Botswana and of Namibia (Research permit CYSC 1/17/2 IV (8) from the Ministry of Youth Sport and Culture of Botswana, and 17/3/3 from the Ministry of Health and Social Services of Namibia). Samples from several populations of southwestern Angola were collected as described by [21]. Each individual gave written consent after the purpose of the study was explained with the help of local translators. Individuals were assigned to populations according to the ethnolinguistic affiliation (i.e., primary language spoken) of their maternal grandmother, as declared during sample collection.

### Samples and mtDNA sequence data

Details on the samples and DNA extraction are given in [12,17,20,21]. Full mtDNA sequence data were generated from genome libraries tagged with either single or double indexes, and enriched for mtDNA following protocols described previously [30,31]. The libraries were sequenced on the Illumina GAIIx (Solexa) platform, using either single or paired end runs of 76 bp length, resulting in an average coverage of ∼400x. Sequences were manually checked with Bioedit (www.mbio.ncsu.edu/BioEdit/bioedit.html) and read alignments were screened with ma [32] to confirm indels. To minimize the impact of missing data, we applied imputation using stringent criteria, replacing missing sites with the nucleotide that was present in at least two otherwise identical haplotypes of the dataset.

One hundred and ninety seven sequences from Botswana, Namibia, and Angola were previously included in studies focusing on haplogroups L0d and L0k as well as on the prehistory of Khoisan populations [18,33], while a subset of 169 sequences from Zambia were included in [20]; the GenBank accession numbers of these samples can be found in Table S1. The remaining 446 sequences from Zambia and 170 sequences from Angola have not yet been reported and are available from GenBank with accession numbers KJ185394-KJ186009. The final alignment consists of 982 sequences of 16465 bp. Positions with missing data as well as the poly-C-regions (np 303-315, 16183-16194) were removed from all analyses; in addition, 45 positions with indels were removed from all analyses run in Arlequin and from sequences used for network construction.

**Table 1:** List of populations included in the study with their linguistic affiliation, geographic location, and values of genetic diversity.

| Population | Country of Sampling | Code | Linguistic Affiliation | Subsistence | N | n | S | seq div | SD | $\pi$ | SD |
| --- | --- | --- | --- | --- | --- | --- | --- | --- | --- | --- | --- |
| Nyaneka | Angola | NYA | West Bantu | agropastoralist | 59 | 50 | 481 | 0.99 | 0.00 | 0.0037 | 1.80E-03 |
| Ovimbundu | Angola | OVM | West Bantu | agriculturalist | 60 | 53 | 494 | 1.00 | 0.00 | 0.0036 | 1.73E-03 |
| Ganguela | Angola | GAN | West Bantu | agriculturalist | 20 | 16 | 290 | 0.97 | 0.03 | 0.0040 | 2.02E-03 |
| Chokwe, Luchazi, Luvale<sup>1</sup> | Zambia | CHO | West Bantu | agriculturalist | 33 | 32 | 319 | 1.00 | 0.01 | 0.0038 | 1.88E-03 |
| Mbunda | Zambia | MBN | West Bantu | agriculturalist | 67 | 55 | 551 | 0.99 | 0.01 | 0.0038 | 1.84E-03 |
| Kuvale | Angola | KUV | West Bantu | pastoralist | 53 | 29 | 369 | 0.95 | 0.02 | 0.0040 | 1.93E-03 |
| Herero | Namibia | HER | West Bantu | pastoralist | 30 | 20 | 160 | 0.94 | 0.03 | 0.0022 | 1.12E-03 |
| Himba | Namibia | HIM | West Bantu | pastoralist | 21 | 13 | 147 | 0.93 | 0.04 | 0.0022 | 1.11E-03 |
| Kgalagadi | Botswana | KGA | East Bantu | agropastoralist | 19 | 15 | 205 | 0.97 | 0.03 | 0.0033 | 1.68E-03 |
| Tswana | Botswana | TSW | East Bantu | agropastoralist | 20 | 19 | 188 | 0.99 | 0.02 | 0.0034 | 1.72E-03 |
| Wider Shona<sup>1</sup> | Botswana | SHO | East Bantu | agriculturalist | 20 | 20 | 317 | 1.00 | 0.02 | 0.0040 | 2.03E-03 |
| Lozi | Zambia | LOZ | East Bantu | agriculturalist | 118 | 94 | 613 | 0.99 | 0.00 | 0.0037 | 1.80E-03 |
| Kwamashi | Zambia | KWA | West Bantu | agriculturalist | 35 | 27 | 319 | 0.97 | 0.02 | 0.0036 | 1.76E-03 |
| Mbukushu | Zambia | MBK | West Bantu | agriculturalist | 23 | 21 | 316 | 0.99 | 0.02 | 0.0038 | 1.91E-03 |
| Shanjo | Zambia | SHA | East Bantu | agriculturalist | 25 | 18 | 299 | 0.97 | 0.02 | 0.0038 | 1.89E-03 |
| Wider Luyana<sup>1</sup> | Zambia | LUY | West Bantu | agriculturalist | 106 | 83 | 545 | 0.99 | 0.00 | 0.0036 | 1.72E-03 |
| Nkoya | Zambia | NKO | West Bantu | agriculturalist | 32 | 25 | 323 | 0.98 | 0.01 | 0.0036 | 1.78E-03 |
| Eastern Tonga<sup>1</sup> | Zambia | TNG | East Bantu | agriculturalist | 48 | 47 | 444 | 1.00 | 0.00 | 0.0038 | 1.86E-03 |
| Fwe | Zambia | FWE | East Bantu | agriculturalist | 33 | 20 | 292 | 0.93 | 0.03 | 0.0038 | 1.87E-03 |
| Subiya | Zambia | SUB | East Bantu | agriculturalist | 20 | 17 | 229 | 0.98 | 0.02 | 0.0035 | 1.78E-03 |
| Totela | Zambia | TOT | East Bantu | agriculturalist | 29 | 25 | 306 | 0.99 | 0.01 | 0.0036 | 1.78E-03 |
| Northeast Zambia<sup>1</sup> | Zambia | NEZ | East Bantu | agriculturalist | 24 | 23 | 334 | 1.00 | 0.01 | 0.0038 | 1.91E-03 |
| Damara | Namibia | DAM | Khoisan | forager/small stock | 38 | 20 | 250 | 0.89 | 0.04 | 0.0025 | 1.23E-03 |

<sup>1</sup>Aggregate populations: ethnolinguistic affiliation of each individual can be traced in Supplementary Table S1.
code = abbreviation used in figures; N = sample size; n = number of haplotypes; S = number of segregating sites; seq div = sequence diversity; SD = standard deviation; $\pi$ = nucleotide diversity.

**Figure 1:**
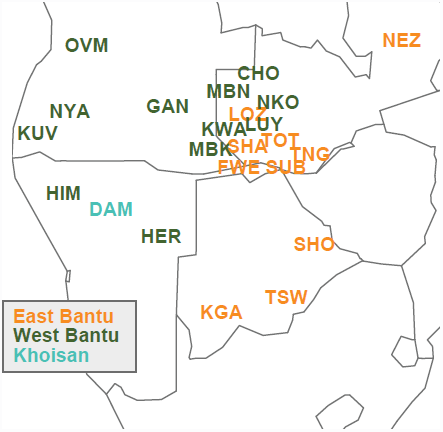
Map showing the rough geographical location of populations, colored by linguistic affiliation. Abbreviations of population labels are as specified in Table 1.

Our dataset includes speakers of several Bantu languages belonging to both the Western and the Eastern branches of Bantu according to the classification found on glottolog 2.2 (http://glottolog.org/). While we were able to group most of the samples into 17 ethnolinguistically homogenous populations that correspond to the identification of donors’ maternal grandmothers, some ethnolinguistic groups were represented by only a few individuals. In these cases, we united samples in five aggregate “populations” of speakers of closely related varieties based mainly on linguistic criteria, but ensuring that the resulting populations were genetically homogeneous as shown by non-significant between-population variance in AMOVA analyses and non-significant ΦST distances (see Table S1 for the ethnolinguistic affiliation of each individual and the composition of the aggregate “populations”). In addition, 49 individuals sampled in Namibia and Zambia could not be grouped into populations with a meaningful sample size or had an unclear ethnic affiliation; these were labeled as “others” and included only in lineage-based analyses (i.e. networks and phylogenetic trees). Supplementary Table S1 provides details on the country of sampling, ethnolinguistic affiliation, and GenBank accession number for each sample. We also included the Damara, who speak a Khoe language rather than a Bantu language, because of their known genetic proximity to Herero and Himba [17,18]. The rough geographic location of the 23 populations included in the study can be seen in Figure 1, while Table 1 summarizes the information on their country of origin, linguistic affiliation, and subsistence.

To investigate the variation in haplogroups L3d and L3f we further included published data from African populations: for the visual depiction in Surfer maps, haplogroup frequency data was collected from the literature as summarized in Table S2, while for networks and BEAST runs, 28 published complete mtDNA genome sequences available from GenBank were added to the alignment [34–38].

### Statistical analyses

Haplogroup assignment was performed with the online tool Haplogrep [39]. Haplogroup affiliation of individuals belonging to haplogroup L0d and L0k was further defined following the nomenclature reported in [33] (see Table S1). Analyses of Molecular Variance (AMOVA), values of sequence diversity, and matrices of pairwise Φ_ST_ values were computed in Arlequin ver. 3.11, while values of nucleotide diversity and variance were calculated in R with the package Pegas [40]. A correspondence analysis (CA) based on haplogroup frequencies was performed with the package ca [41], and non-metric Multi-Dimensional Scaling (MDS) analyses based on pairwise Φ_ST_ values were performed with the function “isoMDS” from the package MASS [42]. The Africa-wide variation in frequency of haplogroups L3d and L3f was visualized on a map with the software Surfer ver. 10.4.799 (Golden Software Inc.). Median-joining networks [43] with all sites given equal weights and no pre- or post-processing steps were computed with Network 4.11 (www.fluxus-engineering.com) and visualized in Network publisher 1.3.0.0. A Mantel test was performed between genetic (Φ_ST_) and geographic distances with the R package vegan [44]; geographic distances between populations were averaged over GPS data from the individual sampling locations with the function rdist.earth of the package fields [45].

BEAST (v1.7.2; [46]) was used to construct Bayesian Skyline Plots and phylogenetic trees, based on the complete mtDNA sequence and using the mutation rate of 1.665 x 10^-8^ from Soares et al. [47]. A Generalized Time Reversible model was applied, and multiple runs were performed for each dataset, using 10, 20 or 30 million chains for single haplogroups and populations. For the schematic tree of the whole dataset 40 million chains and a GTR mutation model were used. The most probable tree from the BEAST runs was assembled with TreeAnnotator and drawn with FigTree v 1.4.0.

Resampling tests were performed in R to investigate the possible shared ancestry of the Herero, Himba, and Damara on the one hand, and the Herero, Himba, and Kuvale on the other, notwithstanding the absence of haplogroup L3f in the Damara and the absence of L3d in the Kuvale and the concomitant high frequencies of these haplogroups in the Herero and Himba. In both cases we proceeded as follows: the Herero and the Himba were considered a single population with a sample size of 51 individuals, while for the Damara and Kuvale we used the actual sample sizes included in the study (i.e. 38 and 53 individuals, respectively). We then created a hypothetical ancestral population with a range of frequencies of the haplogroup of interest. This ancestral population was split into two daughter populations with Ne = 1000, which were resampled for a number of generations proportional to 500, 1000 and 2000 years (with a generation time of 25 years). At the end, a number of individuals corresponding to the population samples of interest (i.e. 53 for Kuvale, 51 for Himba/Herero, or 38 for Damara) were sampled 100 times from the two daughter populations, and the probability of having a frequency of the haplogroup of interest within the range of the respective confidence intervals for both populations was recorded. The whole process was repeated 10,000 times for each initial haplogroup frequency tested, and the probability values were recorded in a table. For all simulations, population size was kept constant and no migration between the daughter populations was considered.

## RESULTS

### Genetic structure of southern African Bantu-speakers

**Figure 2:**
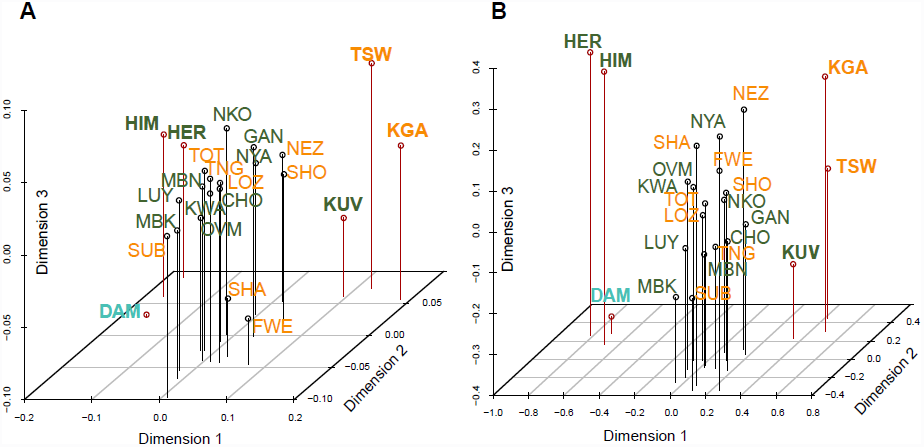
Three-dimensional MOS analysis based on pairwise ΦST values between populations. Color coding by linguistic affiliation; abbreviations of population labels are as specified in Table 1. A. Including all sequences, stress = 5.35 B. Excluding L0d and L0k sequences, stress = 5.34. Populations discussed in the main text are highlighted with bold font and a red line in the plot.

As can be seen from Table S3 and Figure S1, haplogroups found in relatively high frequency across most of the populations of the dataset are L0a, L1c, L2a, and L3e. Other haplogroups, however, are more restricted, being found in only a few populations; of these, L3d and L3f (discussed in detail below) show a particularly striking distribution, being found in very high frequency only in the populations of Namibia.

There is very little discernible structure in the maternal genepool of the Bantu-speaking populations of southern Africa, as shown by a three-dimensional MDS analysis. Only two distinct groups of populations emerge (Figure 2A): the Himba, Herero, and the non-Bantu-speaking Damara from Namibia on the one hand, and the Kuvale from Angola as well as the Tswana and Kgalagadi from Botswana on the other hand; the Tswana and Kgalagadi are separated from their geographic neighbors the Wider Shona. The third dimension, however, splits the Damara from the Himba and Herero. It is notable that the Kuvale are closer to other Bantu-speaking groups than to the Himba and the Herero, who are genetically more similar to the Khoe-speaking Damara in spite of being culturally and linguistically related to the Kuvale.

Haplogroups L0d and L0k are known to be an introgression from autochthonous populations [18,24,33,35] and are thus indicative of post-immigration contact rather than reflecting the genetic relationships among Bantu-speakers themselves; the differential effects of gene flow from autochthonous populations are addressed below. When excluding these clearly introgressed lineages from the MDS analysis, the same two distinctive groups of populations, namely the Kuvale, Tswana, and Kgalagadi, and the Himba, Herero, and Damara, emerge in an even more pronounced manner, with the third dimension again separating the Damara from the Himba and Herero; all the other Bantu-speaking populations, in contrast, cluster very closely (Figure 2B).

As can be seen in the CA plots displayed in Figure S2, the distinct position of the Himba, Herero and Damara populations is driven by their high levels of haplogroup L3d (Figure S2A), which is completely absent from the Kuvale (Table S3). In contrast, the Tswana and Kgalagadi, who stand out in the MDS analysis, are no longer separated in these CA plots, suggesting that their separate position in the MDS plots is mainly due to divergent sequence types rather than a distinct haplogroup composition. The CA plots additionally highlight other aspects of the data, separating the Northeast Zambia population (characterized by the presence of the otherwise largely absent haplogroups L0f and L4) or the Fwe and Shanjo, who have high frequencies of haplogroup L0k (Figure S2A). When excluding these outliers, populations with very high frequencies of haplogroup L0d (Kgalagadi, Tswana, Wider Shona, and Kuvale) stand out (Figure S2B). The influence of different levels of admixture with autochthonous populations on the genetic structure of the southern African Bantu-speaking groups is additionally illustrated by the lack of discernable clusters when the introgressed haplogroups L0d and L0k are excluded (Figure S2C).

The difference of the Himba, Herero, and Damara from the other populations included in this study also becomes apparent from measures of genetic diversity (Table 1): while diversity is high for the Bantu-speaking populations in general, with many of the ethnolinguistically defined self-identified groups (e.g. Nyaneka, Ovimbundu, Kwamashi, Mbukushu) having values of sequence diversity of 0.99-1.00, and with nucleotide diversity ranging between 0.0033 and 0.0040, the Himba, Herero and Damara stand out in having both very low sequence diversity (0.93, 0.94, and 0.89, respectively) and nucleotide diversity (0.0022 for the Herero-speakers, 0.0025 for the Damara). The Kuvale again differ from the other Herero-speakers: although their sequence diversity is relatively low (0.95), their nucleotide diversity is twice as high as that of their linguistic and cultural relatives (0.0040); this diversity pattern resembles that of the Fwe from southwestern Zambia, who have a sequence diversity of only 0.93, but nucleotide diversity of 0.0038.

**Table 2:** Results of AMOVA analyses

|  | **n of groups** | **between groups** | **between pops (within groups)** | **within pops** |
| --- | --- | --- | --- | --- |
| **a) Including haplogroups L0d/L0k** |  |  |  |  |
| All 23 populations | 1 |  | 5.51** | 94.49 |
| 20 populations (excl. HER, HIM, DAM) | 1 |  | 3.22** | 96.78 |
| Linguistic criteria (West vs East Bantu) <sup>a</sup> | 2 | -0.14 | 4.72** | 95.42 |
| Subsistence criteria (Pastoralists vs non-pastoralists) <sup>a</sup> | 2 | 2.64* | 3.97** | 93.39 |
| Geographic Criteria (NW, SW, SE, Centre, NE) <sup>b</sup> | 5 | 4.80** | 1.77** | 93.43 |
| **b) Excluding haplogroups L0d/L0k** |  |  |  |  |
| All 23 populations | 1 |  | 5.94** | 94.06 |
| 20 populations (excl. HER, HIM, DAM) | 1 |  | 2.58** | 97.42 |
| Linguistic criteria (West vs East Bantu) <sup>a</sup> | 2 | -0.33 | 4.85** | 95.48 |
| Subsistence criteria (Pastoralists vs non-pastoralists) <sup>a</sup> | 2 | 3.50* | 3.83** | 92.67 |
| Geographic Criteria (NW, SW, SE, Centre, NE) <sup>b</sup> | 5 | 5.31** | 1.64** | 93.06 |

\* significant at 0.05 level; \*\* significant at 0.01 level
<sup>a</sup> The grouping by linguistic and subsistence criteria followed the assignment in Table 1
<sup>b</sup> Geographic grouping: NW = OVM, NYA, KUV, GAN; SW = HER, HIM; SE = KGA, TSW, SHO; NE = NEZ; CENTRE = CHO, MBN, NKO, LOZ, LUY, KWA, SHA, MBK, TOT, FWE, SUB, TNG
Note: The groupings by linguistic, subsistence, and geographic criteria were performed without the Damara, as these cannot be assigned to the linguistic grouping West Bantu or East Bantu

An Analysis of Molecular Variance (Table 2) demonstrates the relative lack of differentiation among the southern African populations, with only ∼6% of the variation being found between populations, irrespective of whether haplogroups L0d and L0k, which stem from post-immigration admixture, are included or not. A large proportion of the variance between populations is due to the differentiation of the Himba, Herero, and Damara, as shown by the fact that the between-population variance drops to ∼3% when excluding these populations. Affiliation to either of the two major branches of the Bantu family (Eastern Bantu vs. Western Bantu) does not account for any genetic structure, as seen by the complete absence of variance between groups. The three pastoralist populations Herero, Himba, and Kuvale, are somewhat distinct from the non-pastoralist Bantu-speaking populations, as shown by the significant between-group variance of 2.6% and 3.5%, respectively, depending on whether haplogroups L0d and L0k are included in the analysis or not. Nevertheless, the genetic variation of the populations included in the “pastoralist” and “non-pastoralist” grouping is higher than that between the groups. Only a rough geographic subdivision correlates with some degree of genetic structure: in this case the between group variance rises to ∼5% (as opposed to a within group variance of ∼1.7%). On a finer scale, too, the pairwise geographic distances correlate with the genetic distances: a Mantel test gives significant correlations both when including and excluding L0d and L0k sequences (r = 0.3286/p = 0.015 and r = 0.2575/p = 0.043, respectively).

### Haplogroups L0d and L0k

**Figure 3:**
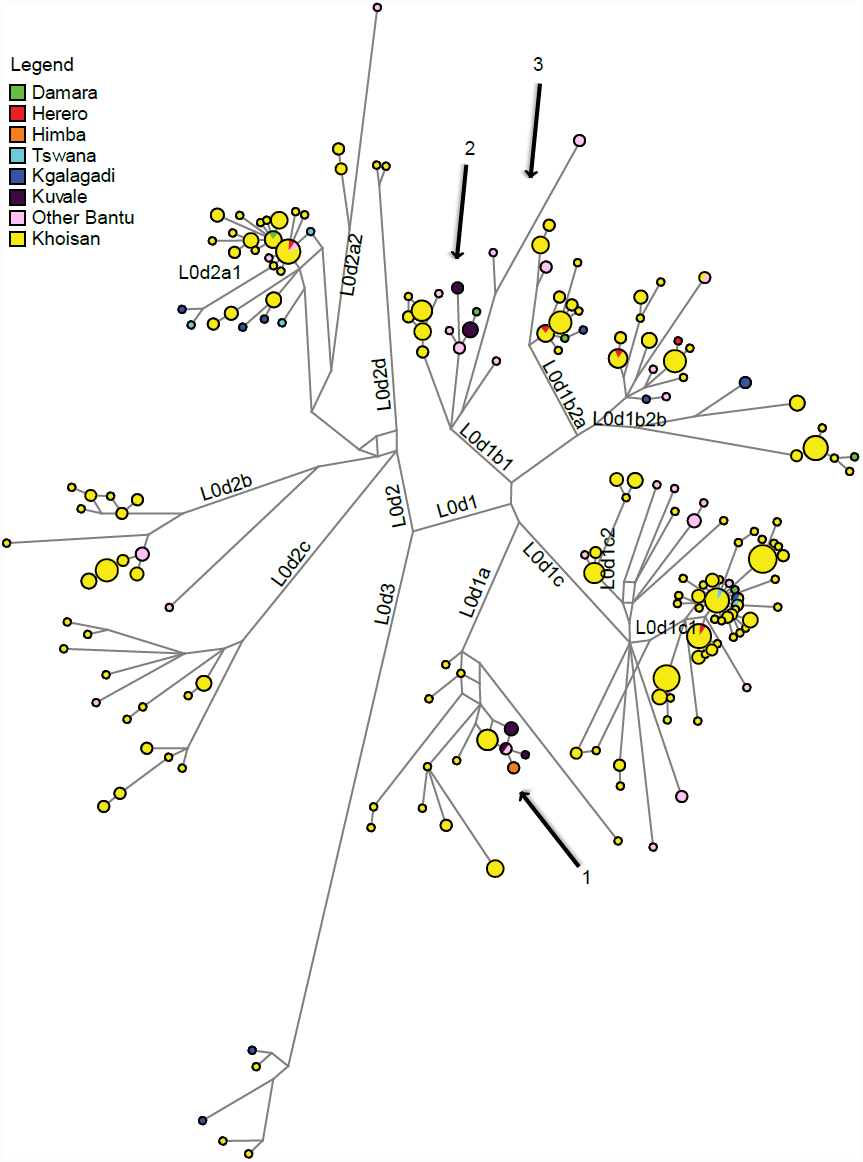
Network of complete mtDNA genome sequences from southern Africa belonging to haplogroup L0d. Branches highlighted by arrows are discussed in the text. Only sublineages of L0d2a1, L0d1b2a, L0d1b2b and L0d1c1 are shared directly between Bantu-speaking and Khoisan-speaking populations.

The mtDNA haplogroups L0d and L0k have been convincingly shown to be characteristic of autochthonous populations of southern Africa [18,24,35]. They therefore represent an ideal measure for detecting admixture in the maternal line between the immigrating Bantu-speaking groups and these autochthonous populations. The frequency of these haplogroups ranges from complete absence in some of the populations of Zambia to 53% in the Kgalagadi of southern Botswana (Table S3, see Figure S3A for a graphic representation of the distribution of L0d and L0k lineages in the populations considered here). Interestingly, hardly any of the L0d and L0k sequences found in the Bantu-speaking populations are directly shared with extant Khoisan foragers or pastoralists. As shown previously [33], the L0k sequences found in Bantu-speaking populations diverge considerably from those found in extant Khoisan populations. With respect to haplogroup L0d, as shown by the network only three Bantu-speaking populations (involving four Herero, two Tswana, and one Kgalagadi individual) share sequences directly with Khoisan (Figure 3). Three branches of the network are found nearly exclusively in Bantu-speaking populations: one of these (belonging to subhaplogroup L0d1a and indicated by arrow 1 in Figure 3) is derived from a sequence type restricted to Khoe-speaking Shua from northeastern Botswana and is found in Kuvale and Himba, with one Kuvale type shared with Nyaneka. The two others (belonging to subhaplogroup L0d1b1 and indicated by arrow 2 and 3 in the figure) are at least 11 mutations distant from the closest Khoisan haplotype; the eight divergent haplotypes found on these branches again belong to Kuvale as well as to different populations of Zambia and Angola. The only population found in this branch that does not speak a Bantu language is the Damara.

### Haplogroups L3d and L3f

The Himba and Herero stand out among the Bantu-speaking populations of southern Africa in having very high frequencies of haplogroups L3d (38% and 47%, respectively) and L3f (29% and 33%, respectively), while in their cultural and linguistic relatives, the Kuvale, L3d is absent and L3f has a frequency of only 6% (Table S3). In contrast, the geographic neighbors of the Himba and Herero, the Khoe-speaking Damara, have 63% L3d but completely lack L3f (cf. Table S3). The high levels of these two haplogroups are thus clearly of key importance for understanding the prehistory of the Herero, Himba, Kuvale, and Damara populations.

**Figure 4:**
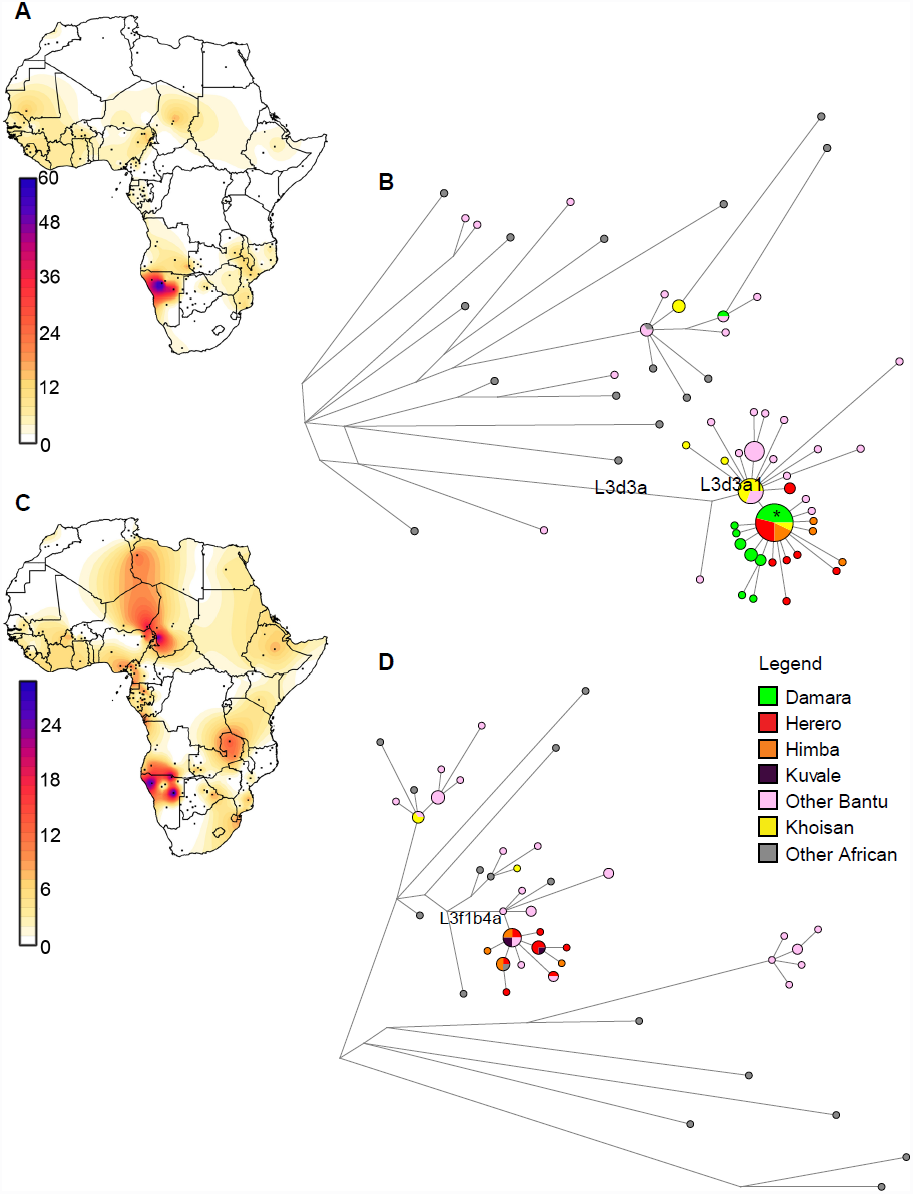
Surfer maps and networks of haplogroups L3d and L3f. A: Surfer map of L3d frequencies in Africa. B: Network of African complete mtDNA genome sequences belonging to haplogroup L3d. C: Surfer map of L3f frequencies in Africa. D: Network of African complete mtDNA genome sequences belonging to haplogroup L3f.

While haplogroup L3d is found across Africa at low frequency (Figure 4A, see Supplementary Table S2 for the populations included in the Surfer map), the lineages found at high frequency in southwestern Africa mostly belong to a single, highly divergent branch, namely L3d3a1 (Figure 4B, Figure S3B). This can be further divided into two clusters: one major node predominating in Khoisan, from which several haplotypes found in Zambian populations radiate, and a derived starlike cluster found mainly in the Himba, Herero, and Damara (indicated by an asterisk in Figure 4B). Dating the signal of expansion detectable in the L3d3a1 branch with the rho statistic [48] and the calculator from Soares et al. [47] gives an age of 395-6668 years BP, while the expansion detectable in the Himba, Herero, and Damara dates to 711-2130 BP. The first date is in good accordance with the pronounced branching dating to 2,500-3,000 years ago in a Bayesian tree of L3d sequences (highlighted in Figure S4A).

In contrast to L3d, L3f is found in frequencies >20% not only in southwestern Africa, but also in some populations of the Cameroon/Chad border areas ([49,50]; Figure 4C, Table S2). Nevertheless, the sequences found in the Himba and Herero all fall onto one restricted branch L3f1b4a (Figure 4D, Figure S3B). Several other Bantu-speaking populations from Namibia and Angola fall into this cluster as well, and the Himba and Herero share two haplotypes with their cultural and linguistic relatives, the Kuvale. This cluster exhibits a signal of expansion which can be dated with the rho statistic to between 526 and 4234 years BP; this corresponds to a pronounced branching 2,500-3,000 years ago in the Bayesian tree of L3f sequences (highlighted in Supplementary Figure S4B).

**Table 3:** Results of resampling tests

| **Initial Percentage** | **0.09** | **0.13** | **0.17** | **0.21** | **0.25** | **0.29** | **0.33** | **0.37** | **0.41** | **0.45** | **0.49** | **0.53** | **0.57** |
| --- | --- | --- | --- | --- | --- | --- | --- | --- | --- | --- | --- | --- | --- |
| 500 ya <sup>a</sup> | 0.02 | 0.11 | 0.32 | 0.47 | 0.62 | 0.65 | 0.51 | 0.47 | 0.22 | 0.11 | 0.08 | 0.01 | 0.00 |
| 1000 ya <sup>a</sup> | 0.04 | 0.11 | 0.26 | 0.40 | 0.55 | 0.58 | 0.52 | 0.39 | 0.18 | 0.11 | 0.09 | 0.03 | 0.01 |
| 2000 ya <sup>a</sup> | 0.03 | 0.12 | 0.24 | 0.31 | 0.41 | 0.42 | 0.36 | 0.30 | 0.22 | 0.20 | 0.10 | 0.05 | 0.02 |
| **Initial Percentage** | **0.01** | **0.05** | **0.09** | **0.13** | **0.17** | **0.21** | **0.25** | **0.29** | **0.32** | **0.34** | **0.36** | **0.38** | **0.4** |
| 500 ya <sup>b</sup> | 0.00 | 0.00 | 0.04 | 0.06 | 0.08 | 0.05 | 0.01 | 0.01 | 0.00 | 0.00 | 0.00 | 0.00 | 0.00 |
| 1000 ya <sup>b</sup> | 0.00 | 0.01 | 0.05 | 0.11 | 0.09 | 0.09 | 0.04 | 0.02 | 0.01 | 0.01 | 0.01 | 0.00 | 0.00 |
| 2000 ya <sup>b</sup> | 0.01 | 0.03 | 0.09 | 0.10 | 0.10 | 0.08 | 0.07 | 0.03 | 0.02 | 0.02 | 0.01 | 0.01 | 0.01 |
| 500 ya <sup>c</sup> | 0.00 | 0.02 | 0.04 | 0.20 | 0.17 | 0.13 | 0.15 | 0.10 | 0.05 | 0.05 | 0.03 | 0.01 | 0.01 |
| 1000 ya <sup>c</sup> | 0.00 | 0.02 | 0.07 | 0.14 | 0.27 | 0.25 | 0.11 | 0.07 | 0.08 | 0.03 | 0.02 | 0.01 | 0.01 |
| 2000 ya <sup>c</sup> | 0.00 | 0.03 | 0.08 | 0.13 | 0.16 | 0.17 | 0.18 | 0.10 | 0.10 | 0.07 | 0.05 | 0.04 | 0.04 |
| 500 ya <sup>d</sup> | 0.00 | 0.00 | 0.00 | 0.02 | 0.03 | 0.01 | 0.01 | 0.00 | 0.00 | 0.00 | 0.00 | 0.00 | 0.00 |
| 1000 ya <sup>d</sup> | 0.00 | 0.00 | 0.01 | 0.01 | 0.02 | 0.01 | 0.01 | 0.01 | 0.00 | 0.00 | 0.00 | 0.00 | 0.00 |
| 2000 ya <sup>d</sup> | 0.00 | 0.01 | 0.02 | 0.03 | 0.03 | 0.02 | 0.02 | 0.01 | 0.01 | 0.01 | 0.01 | 0.00 | 0.00 |

<sup>a</sup> Probability of retaining the shared L3d haplotype in Himba/Herero and Damara for different time splits (Years Ago)
<sup>b</sup> Probability of retaining L3f in Himba and Herero and losing it in Damara for different time splits (Years Ago)
<sup>c</sup> Probability of retaining L3f in Himba and Herero and having it at low percentage in Kuvale for different time splits (Years Ago)
<sup>d</sup> Probability of retaining L3d in Himba and Herero and losing it in Kuvale for different time splits (Years Ago)

The Damara, who have the highest frequency of L3d and who share a highly frequent L3d haplotype with the Herero and Himba, entirely lack L3f. This finding is compatible with two scenarios: 1) The mtDNA pool of the Damara, the Himba, and Herero is derived from a single ancestral population, and haplogroup L3f was lost in the Damara due to genetic drift. 2) The Damara mtDNA lineages stem from a different ancestral population than the Himba and Herero, and the Himba and Herero incorporated large amounts of haplogroup L3d sequences through gene flow from Damara. In order to distinguish between these hypotheses, we performed resampling tests, assuming a frequency of 31% L3f (with 95% confidence intervals (C.I.) 19-46%) in the Himba/Herero as well as 24% of the single L3d3a haplotype shared with the Damara (C.I. 13-37%). For the Damara, a lack of haplogroup L3f (C.I. 0-9%) and 32% of the L3d3a haplotype shared with Himba and Herero (C.I. 18-49%) was assumed. In addition, in the resampling test in which we tried to assess the probability that the Himba, Herero, and Damara would have retained a single shared haplotype at high frequency, we included a probability of change of the L3d3a haplotype each generation with a rate of one mutation every 3533 years following the rate of Soares et al. [47] for the full mtDNA genome. As can be seen from Table 3, the presence of the L3d haplotype shared at high frequency by the Himba, Herero, and Damara is expected with a probability >0.05 even after a split of 2000 years if the frequency of this haplotype in the ancestral population ranged from ∼10-50%. Conversely, if haplogroup L3f was present in the ancestral population at a frequency of ∼5-30%, it could have drifted to high frequency in the Himba and Herero and subsequently been lost in the Damara if the split took place 2000 years ago. Thus, the scenario of shared ancestry of Damara, Himba and Herero with subsequent loss through drift in the Damara of haplogroup L3f cannot be excluded.

It is likewise intriguing that the culturally and linguistically closely related Herero, Himba, and Kuvale have such divergent mtDNA genepools. This might be explained in two ways: 1) these populations stem from a common ancestral population, and differential gene flow led to their strong divergence; 2) these populations have distinct maternal ancestors and their cultural and linguistic relationship is due to a shift in language and culture. These two alternatives were also assessed with a resampling test, assuming a frequency of 31% L3f (with 95% confidence intervals (C.I.) 19-46%) and 43% haplogroup L3d (C.I. 29-58%) in the Himba/Herero and a frequency of 5.7% L3f (C.I. 1-16%) and a lack of haplogroup L3d (C.I. 0-7%) in the Kuvale. As can be seen from Table 3, the hypothesis of a shared ancestor who carried both L3f and L3d is not compatible with the data: even though haplogroup L3f could have drifted to the observed frequencies if its frequency in the ancestral population ranged from ∼8-31%, haplogroup L3d could not have been simultaneously lost from the Kuvale and drifted to the high frequencies currently observed in the Himba and Herero.

## DISCUSSION

### Genetic diversity of Bantu-speaking populations and Western-Eastern Bantu division

Overall, the Bantu-speaking populations of southern Africa are genetically quite homogenous, with a few exceptions such as the Herero and Himba or Tswana and Kgalagadi. While linguistically the populations can be divided into those speaking Western Bantu languages and those speaking Eastern Bantu languages, this division is not detectable in the maternal genepool, with none of the variance among populations corresponding to this linguistic grouping (Table 2). Furthermore, the amount of haplotypes shared between Eastern and Western Bantu speakers does not differ from the amount of haplotypes shared within each linguistic group: 51 of 258 haplotypes are shared among Eastern Bantu speakers, 80 of 381 haplotypes are shared among Western Bantu speakers, and 52 haplotypes are shared between Eastern and Western Bantu speakers. These results are in good accordance with a previous study [11] and support the suggestion that any potential genetic signal of the split between Eastern and Western populations was subsequently erased through admixture [51]. An alternative explanation for the lack of genetic differentiation between the populations speaking West and East Bantu languages is their possibly quite recent split, since East Bantu languages might be an offshoot of West Bantu languages [51]. Furthermore, the so-called Urewe pottery, the first Early Iron Age ceramic tradition of the Great Lakes region, is often linked with the arrival and spread of Bantu speakers in Eastern Africa [1]. The date of its emergence around 2500 years BP [52,53] can thus serve as an approximate starting point of the Eastern Bantu dispersal [54].

Nevertheless, within the homogenous mtDNA landscape of southern African Bantu-speakers some populations do stand out. The most notable outliers are the Herero and Himba (discussed in detail below); apart from these, the Kgalagadi and Tswana as well as Northeast Zambia are separated in the MDS and CA analysis, respectively. The Tswana and Kgalagadi, who speak closely related languages belonging to the Sotho-Tswana group [55], are characterized by very high frequencies of haplogroup L0d, which provides good evidence for extensive admixture in the maternal line with autochthonous populations (Figure S3A). Nevertheless, this high level of indigenous admixture is not the only reason for their distinctiveness, since they remain separate from other populations in the MDS analysis even when haplogroups L0d and L0k are excluded (Figure 2B). That this separation is mainly due to divergent sequence types rather than a distinct haplogroup composition is demonstrated by the fact that they do not stand out in the CA analysis (Figure S2). The Tswana and Kgalagadi speak closely related languages belonging to the homogenous and close-knit Sotho-Tswana group, which is clearly distinct from surrounding language groups [55,56]. Speakers of these languages immigrated from further southeast into what is now Botswana only 500-800 years BP [57]; they would thus have been relatively isolated from the other Bantu-speaking populations included in this study. The admixture of the Kgalagadi and Tswana with autochthonous populations is likely to have taken place to a large extent before their immigration into their current area of settlement while they were still settled further to the southeast. This is evidenced by their complete lack of L0k (which is found in high frequency in Khoisan populations of western Botswana [33]) and by the distinctiveness of most of their L0d lineages, with only one Kgalagadi and two Tswana L0d sequences shared with Khoisan populations from Namibia and Botswana (Figure 3). The Northeast Zambia population shows some affinities with populations further to the northeast. For instance, it is the only population included here to carry haplogroups L0f and L4; for both of these haplogroups an eastern African origin has been suggested [19,58,59]. The presence of these haplogroups highlights the role played by admixture in the diversification of Bantu-speaking populations [59,60].

### Admixture with autochthonous populations

The degree of admixture between the immigrating Bantu-speaking agriculturalists and autochthonous populations is highly variable. On the one hand, some Zambian populations, such as the Nkoya, the Eastern Tonga, or the Totela, carry no autochthonous lineages at all, while on the other hand the Kuvale, Fwe, Tswana, Wider_Shona, and Kgalagadi have 21-53% of haplogroups L0d and/or L0k. There is a noticeable geographical cline in the presence of these two different autochthonous haplogroups (cf. [33]), with L0d being present at high frequency in Bantu-speaking populations from the southern range of our dataset (Botswana, Namibia, and southern Angola), while L0k is practically absent from these populations (Figure S3A). While the frequency distribution of L0d in our Bantu-speaking populations matches that of extant Khoisan populations (where the highest frequencies of L0d are found in populations of South Africa, Botswana, and Namibia), the highest frequencies of L0k in extant Khoisan populations are found in western Botswana and northeastern Namibia [33]. Together with the fact that practically all of the L0k lineages found in Bantu-speaking populations are highly divergent, this distribution of L0k provides good evidence that the Bantu-speaking immigrants into Zambia intermarried with genetically distinct autochthonous populations who are nowadays extinct [20,33].

L0d, too, provides evidence that the gene flow between indigenous populations and immigrating Bantu speech communities involved genetically distinct autochthonous populations. For instance, there are two divergent branches belonging to subhaplogroup L0d1b1 that are practically restricted to Bantu-speaking populations, being found in the Kuvale and Nyaneka of southwestern Angola and in some populations of western Zambia as well as one Damara (see arrows 2 and 3 in Figure 3). In addition, only the Herero, Tswana, and Kgalagadi share L0d haplotypes directly with neighboring Khoisan populations, and this to differing degrees. The admixture between Herero and Khoisan populations is likely to have taken place quite recently, as they share four out of their five L0d sequences (Table S4). In contrast, the Tswana share only two of their five L0d sequences with different Khoisan populations, while two sequences are at least four to five mutational steps distant from any Khoisan haplotypes. Rather surprisingly, the Kgalagadi, who are the Bantu-speaking population with the highest level of autochthonous haplogroups, share only one out of their ten L0d sequences directly with the neighboring Khoe-speaking G|ui; the other haplotypes are between one and at least eight mutational steps distant from the closest Khoisan sequence type (Table S4). These data indicate that the gene flow from autochthonous populations into most of the Bantu-speaking populations included in the dataset took place a long time ago and/or involved Khoisan populations who did not survive into the present. It is furthermore notable that the Mbukushu, who are reported to have been closely associated with Khwe populations, sharing villages and intermarrying with them [61], do not share any sequences with Khwe.

### Relationships of Kuvale, Himba, Herero, and Damara

The most striking results of this study concern the genetic differences between the culturally and linguistically closely related Himba, Herero, and Kuvale on the one hand and the genetic similarity of the Herero and Himba to the culturally and linguistically distinct Damara, on the other. The Himba, Herero, and Damara differ in their maternal lineages from all other populations included here, as shown by the AMOVA results (Table 2) and the MDS and CA analyses (Figure 2 and Figure S2). This is in good accordance with analyses of genomewide SNP data in which the Himba and Damara also stand out as being distinct from other populations speaking Niger-Congo languages [17]. The results of the resampling tests (Table 3) indicate that it is possible that the Himba, Herero, and Damara all derive their mtDNA pool from the same ancestral population (cf. [28]), a suggestion that fits well with the fact that in previous literature both the Damara and the Herero were referred to as Damara, with the specification “Berg Damara” for the former and “Cattle Damara” for the latter [29]. The low diversity values found in these populations (Table 1) as well as the very high frequencies of only one or two haplogroups (L3d and L3f) indicate that they have undergone a severe founder event or bottleneck (cf. [62]), plausibly at the stage of their common ancestor. As suggested by the starlike pattern evident in the networks of haplogroups L3d and L3f (Figure 4B and 4D) and by the profusion of branches highlighted in the Bayesian trees (Figure S4A and S4B), this bottleneck was followed by an expansion in all three populations. Given the age of the expansion of 2,500-3,000 years, this demographic scenario is not compatible with the known recent genocide experienced by the Herero, which took place only 100 years ago [63]. The signal of the genocide-induced recent bottleneck in the Herero might have subsequently been erased through adoption of a Herero identity by Himba.

However, the complete lack of haplogroup L3f in the Damara suggests that they have had a different demographic trajectory from the Himba and Herero – a hypothesis that is further supported by the Bayesian Skyline Plots for these populations (Figure S5): these show a strong signal of recent expansion for the Herero and Himba but not the Damara. Given the difference in life-style between the Himba and Herero on the one hand and the Damara on the other, it is plausible that the stronger signal of expansion detectable in the former is due to their intensive pastoralism, while the foraging and small-stock herding subsistence of the latter did not permit any major increases in population size. Whether the shared ancestral mtDNA pool derives from a pastoralist population, and the Damara subsequently lost their livestock and reverted to foraging, or whether the ancestral population was largely foraging and the Herero and Himba subsequently adopted cattle pastoralism, cannot be elucidated with these data.

Intriguingly, although the Kuvale are linguistically and culturally closely related to the Herero and the Himba, their mtDNA pool is closest to other Bantu-speaking populations of Angola and Zambia, while displaying high levels of introgression of autochthonous L0d lineages. There are two possible explanations for this discrepancy: 1) the Herero, Himba, and Kuvale stem from genetically distinct ancestral populations but converged culturally and linguistically through language/cultural replacement. In this case, the small amount of haplotypes shared by the Kuvale, Herero, and Himba would stem from recent admixture. 2) The Herero, Himba, and Kuvale all stem from a shared ancestor but subsequently diverged through gene replacement, with admixture with now extinct geographically structured autochthonous populations leading to the strong differences in their mtDNA lineages. According to this hypothesis, the Kuvale would have retained most of the ancestral mtDNA genepool, but would have admixed with peoples essentially bearing L0d lineages, while the Himba, Herero and Damara would have undergone large-scale admixture with people bearing haplogroups L3d and L3f (Figure S3). The results of the resampling test (Table 3) demonstrate that if the Himba, Herero, and Kuvale indeed stem from a common maternal ancestral population, this could have carried haplogroup L3f, but not L3d, so that these populations must have undergone both strong genetic drift as well as differential admixture. That the demographic trajectory of the Kuvale differs from that of the Himba and Herero can also be seen in the Bayesian Skyline plot (Figure S5), where the Kuvale, like the Damara, lack the signal of recent expansion detectable for the Himba and Herero.

The L0d lineages in the Kuvale were previously suggested to possibly stem from admixture with the now extinct Angolan Kwadi [21]. These were a pastoralist population who lived within the Kuvale territory and spoke a language related to the Khoe languages, a family that has been suggested to have been brought to southern Africa by a pre-Bantu migration of pastoralists [64]. Since haplogroup L0d is widespread across Khoisan foragers and pastoralists [18], it is difficult to unambiguously assign the Kuvale L0d lineages to a relatively recent pastoralist migration. However, a branch of haplogroup L0d that is restricted to the Kuvale, Himba, and Nyaneka (indicated by arrow 1 in Figure 3) derives from a sequence type found in seven Shua. These are a Khoe-speaking population of northeastern Botswana who are considered possible descendants of the Khoe-Kwadi-speaking pastoralists who would also have been the ancestors of the Kwadi [64]. Since the Shua are settled so far to the east of the Kuvale, direct admixture seems implausible, raising the possibility that these lineages derive from admixture with Kwadi.

In contrast, subhaplogroup L3d3a has a much more confined distribution and is more likely to have been brought to the area by Khoe-speaking pastoralists as previously suggested [18]. This suggestion is in good accordance with the signal of expansion detectable in these lineages ∼2,500-3000 years BP (Figure S4), since archaeological evidence of pastoralism is detectable from ∼2,200 years in the region [16]. Surprisingly, while roughly 50% of the maternal genepool of the Himba, Herero and Damara appear to stem from this putative Khoe admixture, in analyses of genomewide SNP data the Himba and Damara show no affinities with Khoe-speaking populations [17]. However, since the putative incorporation of Khoe maternal lineages might have involved only a few women related in the maternal line, followed by an expansion of this lineage in the Damara, Himba and Herero ancestor, this lineage would have been retained unchanged due to the specific characteristics of mtDNA. In contrast, the signal of relationship with Khoe-speaking populations may have been lost from the autosomal DNA if this single admixture event was followed by several generations of intermarriage with non-Khoe populations. It thus appears likely that the maternal ancestors of the Kuvale, Herero, Himba, and Damara had a haplogroup composition similar to that found in the Kuvale today, albeit with somewhat higher frequencies of L3f, whereas the shared ancestor of the modern-day Herero, Himba, and Damara incorporated Khoe-speaking women carrying haplogroup L3d; subsequently, haplogroup L3f would have drifted to high frequency in the shared pastoralist ancestor of the Himba and Herero while it was lost from the Damara. Detailed Y-chromosomal analyses of the Damara, Himba, Herero and other Bantu-speaking populations of southern Africa are needed to further investigate the prehistory of these groups.

## CONCLUSIONS

In summary, we have been able to show that the maternal genepool of the Bantu-speaking populations of southern Africa is very homogenous. While the linguistic division into Western and Eastern Bantu does not correlate with genetic divergence, the results of the AMOVA and Mantel analyses demonstrate the impact of geography in structuring the genetic variation. Furthermore, there are big differences in the extent of intermarriage between Bantu-speaking agriculturalists and autochthonous peoples, with some populations showing no evidence of gene flow, while others, like the populations of Botswana, carrying substantial proportions of autochthonous lineages. The lack of L0d/k sequences shared between Bantu and Khoisan populations suggests that the admixture undergone by most of the Bantu-speaking immigrants into southern Africa took place soon after their entering the region and partly involved now-extinct autochthonous populations. Lastly, as shown by the results of our resampling tests, a common ancestral population of the Kuvale, Himba, Herero, and Damara cannot be ruled out. Given that these populations currently differ considerably in lifestyle and language, the history of these populations was clearly complex, with language and culture contact as well as genetic admixture playing important roles. Analyses of the Y-chromosomal diversity will shed further light on these processes.

## ACKNOWLEDGMENTS

This study focuses on the prehistory of populations as reflected in their genetic variation. It does not intend to evaluate the self-identification or cultural identity of any group, which consist of much more than just genetic ancestry. We sincerely thank: all the sample donors for their participation in this study and the governments of Botswana, Namibia, and Zambia for supporting our research; Roland Schröder and Madhusudan Reddy Nandineni for assistance with library preparation; Hongyang Xu for help with the imputation process; Ana Duggan for assistance with the tree analysis; and Mingkun Li for assistance with the bioinformatics analysis. This work was funded by the Max Planck Society; CB was supported by the European Research Council ERC-2011-AdG 295733 grant (Langelin); MV was supported by STAB VIDA, Investigação e Serviços em Ciências Biológicas, Lda., and by the Portuguese Ministry for Science, Technology and Higher Education through PhD grant SFRH/BDE/51828/2012.

## References

1. Phillipson DW (2005) African archaeology. Cambridge: Cambridge University Press.

2. Bostoen K (2007) Pots, words and the Bantu problem: On lexical reconstruction and early African history. J Afr Hist 48.

3. Blench R (2006) Archaeology, Language, and the African Past. Rowman Altamira.

4. Nurse D, Philippson G (2003) Towards a historical classification of the Bantu languages. In: Nurse D, Philippson G, editors. The Bantu Languages. London/New York, Routledge. pp. 164–181.

5. Vansina J (1995) New Linguistic Evidence and “The Bantu Expansion.” J Afr Hist 36: 173–195. doi:10.1017/S0021853700034101.

6. Pakendorf B, Bostoen K, de Filippo C (2011) Molecular perspectives on the Bantu expansion: a synthesis. Lang Dyn Chang 1: 50–88.

7. Diamond J, Bellwood P (2003) Farmers and Their Languages: The First Expansions. Science (80-) 300: 597–603.

8. Schadeberg TC (2003) Historical linguistics. In: Nurse D, Philippson G, editors. The Bantu Languages. London: Routeledge. pp. 143–163.

9. Bostoen K, Grollemund R, Koni Muluwa J (2013) Climate-induced vegetation dynamics and the Bantu Expansion: Evidence from Bantu names for pioneer trees (Elaeis guineensis, Canarium schweinfurthii, and Musanga cecropioides). Comptes Rendus Geosci 345: 336–349.

10. Pour NA, Plaster CA, Bradman N (2012) Evidence from Y-chromosome analysis for a late exclusively eastern expansion of the Bantu-speaking people. Eur J Hum Genet 21: 423–429.

11. Alves I, Coelho M, Gignoux C, Damasceno A, Prista A, et al. (2011) Genetic homogeneity across Bantu-speaking groups from Mozambique and Angola challenges early split scenarios between East and West Bantu populations. Hum Biol 83: 13–38.

12. De Filippo C, Barbieri C, Whitten M, Mpoloka SW, Gunnarsdóttir ED, et al. (2011) Y-chromosomal variation in sub-Saharan Africa: insights into the history of Niger-Congo groups. Mol Biol Evol 28: 1255–1269. doi:10.1093/molbev/msq312.

13. Reid A, Sadr K, Hanson-James N (1998) Herding traditions. In: Lane P, Reid A, Segobye A, editors. Ditswa MMung: The Archaeology of Botswana. Gaborone: Pula Press and The Botswana Society. pp. 81–100.

14. Kinahan J (2011) From the beginning: the archaeological evidence. A History of Namibia: From the Beginning to 1990. London: Hurst and Company. pp. 15–43.

15. Mitchell P (2002) The Archaeology of Southern Africa. Cambridge: Cambridge University Press.

16. Pleurdeau D, Imalwa E, Detroit F, Lesur J, Veldman A, et al. (2012) “Of sheep and men”: earliest direct evidence of caprine domestication in southern Africa at leopard cave (Erongo, Namibia). PLoS One 7: e40340.

17. Pickrell JK, Patterson N, Barbieri C, Berthold F, Gerlach L, et al. (2012) The genetic prehistory of southern Africa. Nat Commun 3: 1143. doi:10.1038/ncomms2140.

18. Barbieri C, Güldemann T, Naumann C, Gerlach L, Berthold F, et al. (2014) Unraveling the complex maternal history of Southern African Khoisan populations. Am J Phys Anthropol 153: 435–448.

19. Salas A, Richards M, De la Fe T, Lareu M V, Sobrino B, et al. (2002) The making of the African mtDNA landscape. Am J Hum Genet 71: 1082–1111.

20. Barbieri C, Butthof A, Bostoen K, Pakendorf B (2013) Genetic perspectives on the origin of clicks in Bantu languages from southwestern Zambia. Eur J Hum Genet 21: 430–436.

21. Coelho M, Sequeira F, Luiselli D, Beleza S, Rocha J (2009) On the edge of Bantu expansions: mtDNA, Y chromosome and lactase persistence genetic variation in southwestern Angola. BMC Evol Biol 9: 80.

22. Wood ET, Stover DA, Ehret C, Destro-Bisol G, Spedini G, et al. (2005) Contrasting patterns of Y chromosome and mtDNA variation in Africa: evidence for sex-biased demographic processes. Eur J Hum Genet 13: 867–876.

23. Quintana-Murci L, Harmant C, Quach H, Balanovsky O, Zaporozhchenko V, et al. (2010) Strong Maternal Khoisan Contribution to the South African Coloured Population: A Case of Gender-Biased Admixture. Am J Hum Genet 86: 611–620.

24. Schlebusch CM, Lombard M, Soodyall H (2013) MtDNA control region variation affirms diversity and deep sub-structure in populations from southern Africa. BMC Evol Biol 13: 1–21.

25. De Filippo C, Heyn P, Barham L, Stoneking M, Pakendorf B (2010) Genetic perspectives on forager-farmer interaction in the Luangwa valley of Zambia. Am J Phys Anthr 141: 382–394.

26. Castrì L, Tofanelli S, Garagnani P, Bini C, Fosella X, et al. (2009) mtDNA variability in two Bantu-speaking populations (Shona and Hutu) from Eastern Africa: Implications for peopling and migration patterns in sub-Saharan Africa. Am J Phys Anthropol 140: 302–311.

27. Güldemann T, Stoneking M (2008) A Historical Appraisal of Clicks: A Linguistic and Genetic Population Perspective. Annual Review of Anthropology. Annual Reviews, Vol. 37. pp. 93–109.

28. Soodyall H, Jenkins T (1992) Mitochondrial DNA polymorphisms in Khoisan populations from southern Africa. Ann Hum Genet 56: 315–324.

29. Barnard ACN (1992) Hunters and herders of southern Africa: a comparative ethnography of the Khoisan peoples. Cambridge; New York: Cambridge University Press.

30. Maricic T, Whitten M, Pääbo S (2010) Multiplexed DNA Sequence Capture of Mitochondrial Genomes Using PCR Products. PLoS One 5: e14004–e14004.

31. Meyer M, Kircher M (2010) Illumina sequencing library preparation for highly multiplexed target capture and sequencing. Cold Spring Harb Protoc 2010: pdb– prot5448.

32. Briggs AW, Good JM, Green RE, Krause J, Maricic T, et al. (2009) Targeted retrieval and analysis of five Neandertal mtDNA genomes. Science (80-) 325: 318.

33. Barbieri C, Vicente M, Rocha J, Mpoloka SW, Stoneking M, et al. (2013) Ancient substructure in early mtDNA lineages of southern Africa. Am J Hum Genet 92: 285–292. doi:10.1016/j.ajhg.2012.12.010.

34. Gonder MK, Mortensen HM, Reed FA, de Sousa A, Tishkoff SA (2007) Whole-mtDNA genome sequence analysis of ancient African lineages. Mol Biol Evol 24: 757–768.

35. Behar DM, Villems R, Soodyall H, Blue-Smith J, Pereira L, et al. (2008) The Dawn of Human Matrilineal Diversity. Am J Hum Genet 82: 1130–1140.

36. Batini C, Lopes J, Behar DM, Calafell F, Jorde LB, et al. (2011) Insights into the Demographic History of African Pygmies from Complete Mitochondrial Genomes. Mol Biol Evol 28: 1099–1110.

37. Hartmann A, Thieme M, Nanduri LK, Stempfl T, Moehle C, et al. (2009) Validation of Microarray-Based Resequencing of 93 Worldwide Mitochondrial Genomes. Hum Mutat 30: 115–122.

38. Torroni A, Achilli A, Macaulay V, Richards M, Bandelt H-J (2006) Harvesting the fruit of the human mtDNA tree. TRENDS Genet 22: 339–345.

39. Kloss-Brandstätter A, Pacher D, Schönherr S, Weissensteiner H, Binna R, et al. (2011) HaploGrep: a fast and reliable algorithm for automatic classification of mitochondrial DNA haplogroups. Hum Mutat 32: 25–32.

40. Paradis E (2010) pegas: an R package for population genetics with an integrated– modular approach. Bioinformatics 26: 419.

41. Nenadic O, Greenacre M (2007) Correspondence analysis in R, with two-and three-dimensional graphics: the ca package. J Stat Softw 20: 1–13.

42. Venables WN, Ripley BD (2002) MASS: modern applied statistics with S. New York: Springer.

43. Bandelt HJ, Forster P, Rohl A (1999) Median-joining networks for inferring intraspecific phylogenies. Mol Biol Evol 16: 37–48.

44. Oksanen J, Blanchet FG, Kindt R, Legendre P, Minchin PR, et al. (2012) vegan: Community Ecology Package. R package version 2.0-5. http://cran.r-project.org/web/packages/vegan/index.html.

45. Furrer R, Nychka D, Sain S (2012) fields: Tools for spatial data. R Packag version 6: http://CRAN.R–project.org/package=fields.

46. Drummond AJ, Suchard MA, Xie D, Rambaut A (2012) Bayesian Phylogenetics with BEAUti and the BEAST 1.7. Mol Biol Evol 29: 1969–1973.

47. Soares P, Ermini L, Thomson N, Mormina M, Rito T, et al. (2009) Correcting for purifying selection: an improved human mitochondrial molecular clock. Am J Hum Genet 84: 740–759.

48. Forster P, Harding R, Torroni A, Bandelt HJ (1996) Origin and evolution of Native American mtDNA variation: a reappraisal. Am J Hum Genet 59: 935–945.

49. Černỳ V, Salas A, Hajek M, Žaloudková M, Brdička R (2007) A Bidirectional Corridor in the Sahel-Sudan Belt and the Distinctive Features of the Chad Basin Populations: A History Revealed by the Mitochondrial DNA Genome. Ann Hum Genet 71: 433–452.

50. Cerezo M, Černỳ V, Carracedo Á, Salas A (2011) New insights into the Lake Chad Basin population structure revealed by high-throughput genotyping of mitochondrial DNA coding SNPs. PLoS One 6.

51. De Filippo C, Bostoen K, Stoneking M, Pakendorf B (2012) Bringing together linguistic and genetic evidence to test the Bantu expansion. Proc R Soc B-Biological Sci 279: 3256–3263.

52. Clist B (1987) A critical reappraisal of the chronological framework of the early Urewe Iron Age industry. Muntu 6: 35–62.

53. Ashley CZ (2010) Towards a socialised archaeology of ceramics in Great Lakes Africa. African Archaeol Rev 27: 135–163.

54. Neumann K, Bostoen K, Höhn A, Kahlheber S, Ngomanda A, et al. (2012) First farmers in the Central African rainforest: A view from southern Cameroon. Quat Int 249: 53–62. doi:10.1016/j.quaint.2011.03.024.

55. Gowlett DF (2003) The Bantu Languages. In: Nurse D, Philippson G, editors. Routledge. pp. 609–638.

56. Creissels D (1999) Remarks on the Sound Correspondences between Proto-Bantu and Tswana (S. 31), with Particular Attention to Problems Involving* j (or* y),* i and sequences* NC. In: Hombert J-M, Hyman LM, editors. Bantu Historical Linguistics. Theoretical and Empirical Perspectives. Stanford: CSLI (Center for the Study of Language and Information) Publications. pp. 297–334.

57. Solway JS (1991) Tswana. In: Middleton J, Rassam A, editors. Encyclopedia of World Cultures, Volume IX - Africa and the Middle East. Boston, G.K. Hall & Company. pp. 360–365.

58. Kivisild T, Reidla M, Metspalu E, Rosa A, Brehm A, et al. (2004) Ethiopian mitochondrial DNA heritage: tracking gene flow across and around the gate of tears. Am J Hum Genet 75: 752–770.

59. Batai K, Babrowski KB, Arroyo JP, Kusimba CM, Williams SR (2013) Mitochondrial DNA Diversity in Two Ethnic Groups in Southeastern Kenya: Perspectives from the Northeastern Periphery of the Bantu Expansion. Am J Phys Anthropol 150: 482–491.

60. Sikora M, Laayouni H, Calafell F, Comas D, Bertranpetit J (2010) A genomic analysis identifies a novel component in the genetic structure of sub-Saharan African populations. Eur J Hum Genet 19: 84–88.

61. Fisch M (2005) The Mbukushu in Angola (1730-2002): A History of Migration, Flight and Royal Rainmaking. Rüdiger Köppe.

62. Spurdle AB, Hammer MF, Jenkins T (1994) The Y Alu polymorphism in southern African populations and its relationship to other Y-specific polymorphisms. Am J Hum Genet 54: 319–330.

63. Wallace M (2011) A History of Namibia: From the Beginning to 1990. Columbia University Press.

64. Güldemann T (2008) A linguist’s view: Khoe-Kwadi speakers as the earliest food-producers of southern Africa. South African Humanit 20: 93–132.

